# Eukaryotic MAGs recovered from deep metagenomic sequencing of the seagrass, *Zostera marina,* include a novel chytrid in the order Lobulomycetales

**DOI:** 10.1101/2025.02.11.637735

**Authors:** Cassandra L. Ettinger, Jonathan A. Eisen, Jason E. Stajich

**Author notes:** **Corresponding Author:** Cassandra L. Ettinger.

## Abstract

Fungi play pivotal roles in terrestrial ecosystems as decomposers, pathogens, and endophytes, yet their significance in marine environments is often understudied. Seagrasses, as globally distributed marine flowering plants, have critical ecological functions, but knowledge about their associated fungal communities remains relatively limited. Previous amplicon surveys of the fungal community associated with the seagrass, *Zostera marina* have revealed an abundance of potentially novel chytrids. In this study, we employed deep metagenomic sequencing to extract metagenome-assembled genomes (MAGs) from these chytrids and other microbial eukaryotes associated with *Z. marina* leaves. Our efforts resulted in the recovery of five eukaryotic MAGs, including a single fungal MAG in the order Loubulomycetales (65% BUSCO completeness), three MAGs representing diatoms in the family Bacillariaceae (93%, 70% and 31% BUSCO completeness) and a single MAG representing a haptophyte algae in the genus *Prymnesium* (40% BUSCO completeness). Whole-genome phylogenomic assessment of these MAGs suggests they all largely represent under sequenced, and possibly novel eukaryotic lineages. Of particular interest, the chytrid MAG was placed within the order Lobulomycetales, consistent with the identity of the dominant chytrid from previous *Z. marina* amplicon survey results. Annotation of this MAG yielded 5,650 gene models of which 77% shared homology to current databases. With-in these gene models, we predicted 121 carbohydrate-active enzymes and 393 secreted proteins (103 cytoplasmic effectors, 30 apoplastic effectors). Exploration of orthologs between the Lobulomycetales MAG and existing Chytridiomycota genomes have revealed a landscape of high-copy gene families related to host recognition and interaction. Further machine learning analyses based on carbohydrate-active enzyme composition predict that this MAG is a symbiont. Overall, these five eukaryotic MAGs represent substantial genomic novelty and valuable community resources, contributing to a deeper understanding of the roles of fungi and other microbial eukaryotes in the larger seagrass ecosystem.

## Introduction

Fungi are key contributors in terrestrial ecosystems, functioning as decomposers, pathogens, and endophytes, but their importance in marine environments remains underexplored [1–3]. While over 120,000 species of terrestrial fungi have been documented, only about 2,100 marine fungal species have been formally described, though the actual diversity is believed to be much higher [4–7]. Recent research has shed light on the distribution, activity, and biomass of marine fungi, with current estimates suggesting that fungi contribute 0.19-0.21 Gt C to the biomass of the open ocean [8–14]. However, host-associated fungal communities in marine ecosystems, including those associated with marine plants such as seagrasses, are still largely understudied.

Seagrasses are fully submerged flowering plants and vital foundation species in coastal ecosystems globally. *Zostera marina*, commonly known as eelgrass, is a widely distributed species that plays a crucial role in providing essential ecosystem services across coastal regions in the Northern Hemisphere [15–17]. Seagrasses like *Zostera marina* are home to a diverse range of microbial eukaryotes, including diatoms, dinoflagellates, algae, oomycetes, protists, and fungi [18–24]. Studies have shown that fungi represent only a small fraction of the total epiphytic eukaryotic community associated with *Z. marina*, with diatoms, oomycetes, and dinoflagellates among the dominant groups [25–27].

Although culture-based studies have explored fungal diversity associated with *Z. marina*, these approaches are limited by the challenges of growing marine fungi in laboratory settings [28–32]. Advances in DNA-based, culture-independent sequencing techniques have allowed for a more comprehensive characterization of seagrass-associated fungal communities, including their community assembly and biogeographical patterns [25,33–39]. Despite this progress, many seagrass-associated fungi, particularly those from Chytridiomycota lineages, remain difficult to taxonomically classify due to biases in reference databases, which are heavily skewed toward terrestrial fungi [25,36,38,39]. This is unsurprising as a limited number of lineages of marine Chytridiomycota have been described through culture-based methods, and few have sequenced genomes [6,7,40,41]. Nevertheless, marine Chytridiomycota are consistently recovered in DNA-based surveys and are often dominant members of seagrass-associated fungal communities [14,25,34,36,42].

Chytridiomycota in the order Lobulomycetales have previously been observed in high abundance on and inside *Z. marina* leaves [36], and members of this group are part of the *Z. marina* core mycobiome [25]. The Loubulomycetales are a relatively understudied group of chytrid fungi that have been found to span terrestrial, freshwater and marine ecosystems as both parasites and saprobes [43]. The two sequenced genomes from this group represent *Lobulomyces angularis,* which was isolated from an acidic freshwater lake [44], and *Clydaea vesicula,* which was isolated from soil underneath Eucalyptus trees [43]. During the initial description of the Loubulomycetales, the authors noted that due to the difficulty in isolating members of this group, even for ‘chytrid specialists’, that they expected most members would need to be identified and described from molecular data [43].

While previous work identified high relative abundance of chytrid fungi associated with seagrasses, particularly members of the Lobulomycetales, we still have little understanding of the functional role of these enigmatic aquatic chytrids [25,36,45]. To address this, we used a metagenomic sequencing approach on DNA that was previously extracted from *Zostera marina* leaves in Ettinger and Eisen [36] and was reported to have a high relative abundance of Lobulomycetales chytrid fungi. We then used this metagenomic sequencing data to: (i) generate a set of refined eukaryotic metagenome-assembled genomes (MAGs), (ii) phylogenetically place in the tree of life and functionally annotate these eukaryotic MAGs, and (iii) use comparative genomics approaches to explore hypotheses of the possible function and ecology of any fungal MAGs.

## Methods

### Sequence generation

DNA was extracted from epiphytic washes from *Z. marina* leaves as part of previous work focused on characterizing the mycobiome using high-throughput sequencing of the ITS2 region [36]. Three DNA extracts from that work were selected based on predicted relative abundances of an unclassified chytrid in the order Lobulomycetales (identified as amplicon sequence variant SV8), with two samples (108A and 109A) having relatively high abundance of this putative taxa, and one sample having a low abundance of this taxa (140A). These samples with variable abundance of SV8 were chosen to aid binning algorithms which can utilize differential coverage to aid in metagenomic bin recovery [46,47]. DNA was provided to the UC Davis Genome Center DNA Technologies Core for library preparation and sequencing. DNA libraries were sequenced on an Illumina HiSeq4000 to generate 150 bp paired-end reads.

### Sequence processing

Sequence reads were trimmed using bbDuk v. 37.68 [48] with the following parameters: qtrim=rl trimq=10 maq=10. We then mapped against and removed any reads from the metagenomes matching the available genome for *Z. marina* v. 3.1 [49] using bowtie2 v. 2.4.5 [50] and samtools v. 1.11 [51]. The remaining reads from all three metagenomic samples were co-assembled using MEGAHIT v. 1.2.9 [52]. We ran Whokaryote v. 1.1.2 [53], which uses a random forest classifier to predict the eukaryotic origin of contigs. Whokaryote was run with the parameter --minsize 1 kbp and we chose to include predictions from Tiara v. 1.0.3 [53,54], which uses deep-learning to further identify eukaryotic contigs.

The anvi’o v. 7.2 workflow [55] was used to identify and assess eukaryotic MAGs. This workflow was executed twice: once for the entire co-assembly and once for the subset of the co-assembly containing only the Whokaryote predicted eukaryotic contigs. Briefly, this workflow involved obtaining read coverage for each metagenomic sample against the co-assembly using bowtie2 v. 2.4.5 [50] and samtools v. 1.11 [51]. Next “anvi-gen-contigs-database” was used to generate a contig database from the co-assembly and gene calls in the co-assembly were predicted using Prodigal v. 2.6.3 [56]. Single-copy bacterial [57], archaeal [57], and protista [58] genes were then predicted using HMMER v. 3.2.1 [59] and ribosomal RNA genes were predicted using barrnap [60]. Each gene call was then assigned a putative taxonomy using Kaiju v. 1.8.2 [61] with the NCBI BLAST non redundant protein database nr including fungi and microbial eukaryotes v. 2020-05-25. For each metagenomic sample, an individual anvi’o profile was constructed using “anvi-profile” for contigs >1 kbp with the “–cluster-contigs” option. Several automatic binning algorithms were then employed to generate a preliminary set of MAGs including MetaBAT2 v. 2.15, MaxBin v. 2.2.1, BinSanity v.0.5.4 and CONCOCT v. 1.1.0 [62–65]. All predicted MAG bins were then assessed with eukCC v. 2.0 [66] for eukaryotic completeness and redundancy.

To obtain a final representative set of eukaryotic MAGs, dereplication was performed using dRep v. 3.2.2 [67] by providing the eukCC completeness estimates. This step involved combining bins identified from both anvi’o workflow runs (the results from the entire co-assembly and the subset of eukaryotic contigs). The minimum completeness threshold for dRep was set to 1 to ensure a comprehensive representation of eukaryotic MAGs, which often have lower completeness compared to bacterial MAGs.

Representative MAGs with >30% eukaryotic completeness were assessed using anvi-refine. A completeness threshold of >30% has been previously used as a cut-off for high-quality eukaryotic draft MAGs [68]. Further assessment and decontamination of these eukaryotic MAGs was performed using the BlobTools2 v. 4.1.4 [69] workflow. This workflow involves assigning taxonomy based on searches against the UniProt Reference Proteomes database (v. 2021_04) to each contig using diamond (v. 2.1.0) and against the nt database (downloaded 2023-03-21) using command-line BLAST v. 2.2.30+ [70,71]. Next, coverage of each individual metagenomic sample against each eukaryotic bin was calculated with Minimap2 v. 2.24 [72] and sorted using samtools v. 1.11 [51]. Finally the BlobToolKit Viewer was used to visualize the GC content, read coverage and predicted taxonomies of contigs to identify contaminants. To be conservative, any contigs that were taxonomically identified as viral, bacterial, or archaeal were removed from the eukaryotic bins.

After removing potential contamination, we ran EUKulele v. 1.0.2 [73] to assess and refine the initial taxonomic assignments for each bin. We also used BUSCO v. 5.3.2 [74] to assess completion of the de-contaminated bins in ‘genome’ mode using the eukaryotic_odb10 set, and as appropriate the fungi_odb10 or stramenopiles_odb10 sets.

### Annotation

Draft MAG assemblies were screened for contaminant vectors using the ‘vecscreen’ option in the Automatic Assembly of the Fungi (AAFTF) pipeline v. 0.4.1 [75]. Repetitive regions were *de novo* identified and soft masked prior to MAG annotation using RepeatModeler v. 2.0.1 [76] and RepeatMasker v. 4-1-1 [77].

Draft eukaryotic MAGs were annotated using the Funannotate pipeline v. 1.8.14 [78]. Funannotate incorporates multiple software algorithms for eukaryotic gene prediction including Augustus v. 3.4.0, GeneMark-ETS v. 4.71, GlimmerHMM v. 3.0.4, and SNAP v. 2013_11_29 [79–83], and then generates consensus gene models using EVidenceModeler v. 1.1.1 [84]. For two MAGs (SGEUK-04 and SGEUK-05) where GeneMark-ETS was unable to run due to assembly fragmentation, we used –weights genemark:0 during prediction to skip use of this software. Funannotate also uses tRNAscan v. 2.0.11 to predict tRNAs [85]. Annotation of consensus gene models by Funannotate is based on searches against Pfam v. 35.0 [86] and dbCAN v. 9.0 [87,88] using HMMER v.3 [59] and against MEROPS v. 12.0 [89], eggNOG v. 2.1.9 [90], InterProScan v. 5.51-85.0 [91], and UniProt v. 2022_05 [92] using diamond v. 2.0.15 [93]. Transmembrane proteins are further predicted using Phobius v. 1.01 [94] and secreted proteins using SignalP v. 5.0b [95]. For the fungal MAG (SGEUK-03), AntiSMASH v. 6.1.1 was run to predict biosynthetic gene clusters [96] and EffectorP v. 3.0 was run on putative secreted proteins to predict apoplastic and cytoplasmic effectors [97]. We also used CATAStrophy v. 0.1.0 [98], a classification method based on carbohydrate-active enzyme (CAZyme) patterns from filamentous fungal plant pathogens, to predict the possible trophic strategy of SGEUK-03. CATAStrophy was run in a Google Collab implementation using dbCAN v. 10 [87,88].

BUSCO v. 5.3.2 [74] was then used to assess completion of MAG annotations in ‘protein’ mode using the eukaryotic_odb10 set, and as appropriate the fungi_odb10 or stramenopiles_odb10 sets.

### Phylogenetic assessment

To examine the phylogenetic placement of the five eukaryotic MAGs, we carried out analysis of three groups: stramenopiles (SGEUK-01, SGEUK-02, SGEUK-04), fungi (SGEUK-03), and haptista (SGEUK-05). For each group, we obtained gene models from annotated genomes that were publicly available in NCBI GenBank (Tables S1–S3). Given the limited availability of annotated genomes for haptista (four genomes at the time of this work), we also incorporated publicly available transcriptomic data from METdb [99–101].

For stramenopiles, the final dataset for phylogenomic analyses included SGEUK-01, SGEUK-02, SGEUK-04, and 153 annotated genomes representing the following groups: Oomycota (n=120), Bacillariophyta (n=11), Bigyra (n=8), Pelagophyceae (n=7), Eustigmatophyceae (n=3), Phaeophyceae (n=2), Synurophyceae (n=1), and Xanthophyceae (n=1) (Table S1) [102–164]. For fungi, the final dataset for comparative genomics and phylogenomic placement included SGEUK-03 and 98 annotated genomes from the following phyla: Chytridiomycota (n=71), Blastocladiomycota (n=10), Neocallimastigomycota (n=8), Ascomycota (n=4), Monoblepharidomycota (n=3), Olpidiomycota (n=1), and Basidiomycota (n=1) (Table S2) [165–186]. For haptista, the final dataset for phylogenomic placement included SGEUK-05, 47 annotated genomes and transcriptomes from members of the orders Phaeocystales (n=14), Isochrysidales (n=14), Prymnesiales (n=9), Pavlovales (n=7), Coccolithales (n=2), Coccosphaerales (n=1), and Haptophyta *incertae sedis* (n=1) (Table S3) [101,162,187–189].

To place MAGs in a phylogenetic context, the PHYling_unified (https://github.com/stajichlab/PHYling_unified) pipeline was used to generate a protein alignment of each MAG to close relatives based on preliminary taxonomic assignment (i.e., the datasets described in Table S1-S3). This pipeline utilizes HMMER v.3 [59] and ClipKIT [190] to search for, build, and trim an alignment based on gene set. Here we used the most relative BUSCO gene set for each dataset, e.g. the BUSCO fungi_odb10 for SGEUK-3, eukaryotic_odb10 for SGEUK-05, and stramenopiles_odb10 for SGEUK-01, SGEUK-02, and SGEUK-04. Maximum likelihood phylogenies were built from these alignments using IQ-TREE2 v.2.2.6 [191], with the -p option to indicate gene partitions [192] and the -m MFP option to run ModelFinder Plus to identify and use the optimal evolutionary model for each partition [193]. The resulting phylogenetic trees were imported into R v. 4.3.0 [194] and visualized using ggtree v. 3.8.2 [195].

Given that 18S rRNA sequences often do not end up in MAG bins [196] and to further refine taxonomy for the fungal MAG (SGEUK-03), we used phyloflash v. 3.4 [197] to assemble and identify the 18S rRNA gene sequences in the metagenomes. We then identified closely related sequences to the 18S rRNA gene sequence for the fungal MAG using NCBI’s Standard Nucleotide BLAST with default settings [71]. These results were then used to guide a literature search to identify additional sequences for inclusion during phylogenetic reconstruction (Table S4) [43,198–209]. These 18S rRNA gene sequences were then aligned using SSU-ALIGN v. 0.1.1 [210] and trimmed using trimAl v. 1.4 with the -gappyout method [211]. A maximum likelihood phylogeny was built from this alignment using IQ-TREE2 v.2.2.6 [191], with the -p option to indicate gene partitions [192] and the -m MFP option to run ModelFinder Plus to identify and use the optimal evolutionary model for each partition [193]. The resulting phylogenetic tree was imported into R and visualized using ggtree v. 3.8.2 [195].

### Comparative genomics of chytrid MAG

OrthoFinder v. 2.5.4 [212] was used to predict phylogenetic hierarchical orthogroups (HOGs), or gene families, across Chytridiomycota and related lineages (Table S1). Given the little genomic information available for close relatives of SGEUK-03 we focused our analyses on HOGs at the root node (i.e., all HOGs). Due to the incomplete nature of MAGs, we chose to focus comparative analyses on high-copy HOGs, and did not explore gene loss. High-copy HOGs (≥5 copies in SGEUK-03) were visualized in R using the pheatmap v. 1.0.12 package [213].

## Results

### Successful recovery and preliminary annotation of five eukaryotic MAGs

In total, we recovered five eukaryotic MAGs that were high-quality (>30% complete). Briefly, we note that bins resulting from the co-assembly containing only Whokaryote identified eukaryotic contigs had lower contamination, though at the expense of overall completion, compared to their entire co-assembly counterparts. Among the five dereplicated MAGs, only one MAG (SGEUK-05) was not obtained from the eukaryotic co-assembly.

To perform preliminary taxonomic assignment of MAGs, we used EUKulele which reports the relative proportion of all the proteins within a MAG that have annotations that agree at a particular taxonomic level. Using a conservative cut-off of >70% proportional agreement, EUKulele predicted that we had three Bacillariophyta MAGs (SGEUK-01: 89.60%, SGEUK-02: 78.39%, SGEUK-04: 75.08%), one Eukaryota MAG (SGEUK-03: 94.24%), and one Prymnesiophyceae MAG (SGEUK-05: 82.24%).

These eukaryotic draft MAGs were fairly fragmented (i.e., with total contigs ranging from 1,074 to 11,398) likely due to their relatively low mean coverage (i.e., ranging from 3.47x to 7.71x). Briefly, we discuss the general characteristics of the assembly and annotation of each draft MAG (Table 1):

- SG-EUK01 (Bacillariophyta) spans 32.54 Mbp with 3.46x coverage, distributed across 2,172 contigs with an N50 of 20,103 bp. Initial EukCC completion and contamination estimates were 84.5% and 5.7%, respectively. After removing 21 contigs flagged as bacterial contamination by BlobTools, BUSCO analysis using the stramenopiles_odb10 dataset indicated a high completeness of 93.0%. Annotation with Funannotate resulted in 13,603 gene models, including 13,520 mRNA genes and 83 tRNA genes, with 75.92% of gene models having EggNog database hits. BUSCO protein estimates were consistent with the genome completeness, showing 91.0% of the expected single-copy orthologs present using the stramenopiles_odb10 set.
- SG-EUK02 (Bacillariophyta) is 28.49 Mbp in length with 5.37x coverage, spanning 5,479 contigs and an N50 of 5,715 bp. EukCC estimated 64.39% completeness and 13.96% contamination, lower in completeness but higher in contamination than SG-EUK01. We subsequently removed 92 contigs identified as bacterial contamination using BlobTools. BUSCO estimates show 70.0% completeness using the stramenopiles_odb10 dataset. Annotation resulted in 10,889 gene models, including 10,868 mRNA genes and 21 tRNA genes, with 76.43% of gene models having EggNog database hits. BUSCO protein estimates are aligned with genome completeness, showing 69.0% of single-copy orthologs present using the stramenopiles_odb10 set.
- SG-EUK03 (Eukaryota) is a smaller MAG at 12.11 Mbp with 7.71x coverage, distributed across 1,074 contigs with an N50 of 15,102 bp. EukCC predicted completion and contamination at 56.96% and 1.74%, respectively. After removing 10 bacterial contigs identified by BlobTools, BUSCO analysis revealed moderate completeness, with 65.1% using the eukaryota_odb10 dataset and 51.6% using the fungi_odb10 dataset. Annotation resulted in 5,650 gene models, including 5,630 mRNA genes and 20 tRNA genes, with 76.58% of gene models having EggNog database annotation hits. Notably, 393 genes (6.96%) were predicted to be secreted, including 133 potential effectors (103 cytoplasmic, 30 apoplastic), and 121 genes (2.14%) had CAZyme domains including 65 glycosyltransferases (GTs), 26 glycoside hydrolases (GHs), 13 carbohydrate esterases (CEs), 8 polysaccharide lyases (PLs), 7 auxiliary activities (AAs), and 2 carbohydrate-binding modules (CBMs) (Figure S1, Table S5). Based on CAZyme content, CATAStrophy predicted that SGEUK-03 was most likely a symbiont. BUSCO protein estimates indicated 65.9% completeness with eukaryota_odb10 and 57.0% with fungi_odb10, outperforming the genome completeness estimates.
- SG-EUK04 (Bacillariophyta) spans 19.6 Mbp with 4.04x coverage, spread over 11,398 contigs with an N50 of 1,790 bp. EukCC estimated a relatively low completeness of 35.9% and contamination of 5.98%, lower in completeness compared to SG-EUK01 and SG-EUK02. We removed 197 contigs identified as potential bacterial contaminants. BUSCO estimates using the stramenopiles_odb10 dataset indicated that only 31.0% of single-copy orthologs were present and complete. Annotation resulted in 8,359 gene models, including 8,343 mRNA genes and 16 tRNA genes, with 64.77% of gene models having EggNog database hits. BUSCO protein estimates are aligned with the genome completeness, with 30.0% of expected single-copy orthologs present using the stramenopiles_odb10 set.
- SG-EUK05 (Prymnesiophyceae) is the largest of the MAGs, spanning 59.4 Mbp with 5.12x coverage, distributed across 9,735 contigs with an N50 of 6,756 bp. Initial EukCC estimates placed completion at 34.0% and contamination at 3.4%. Following the removal of 197 bacterial contigs, BUSCO estimates showed that 40.0% of expected orthologs were present using the eukaryota_odb10 dataset. Annotation resulted in 15,098 gene models, including 15,075 mRNA genes and 23 tRNA genes, with 56.78% of gene models having EggNog database annotation hits. Repeats represented 18.63% of the genome length, much higher than for any of the other draft MAGs. BUSCO protein estimates performed slightly worse than the genome completeness with only 34.5% of the expected single-copy orthologs present and complete.

**Table 1.**
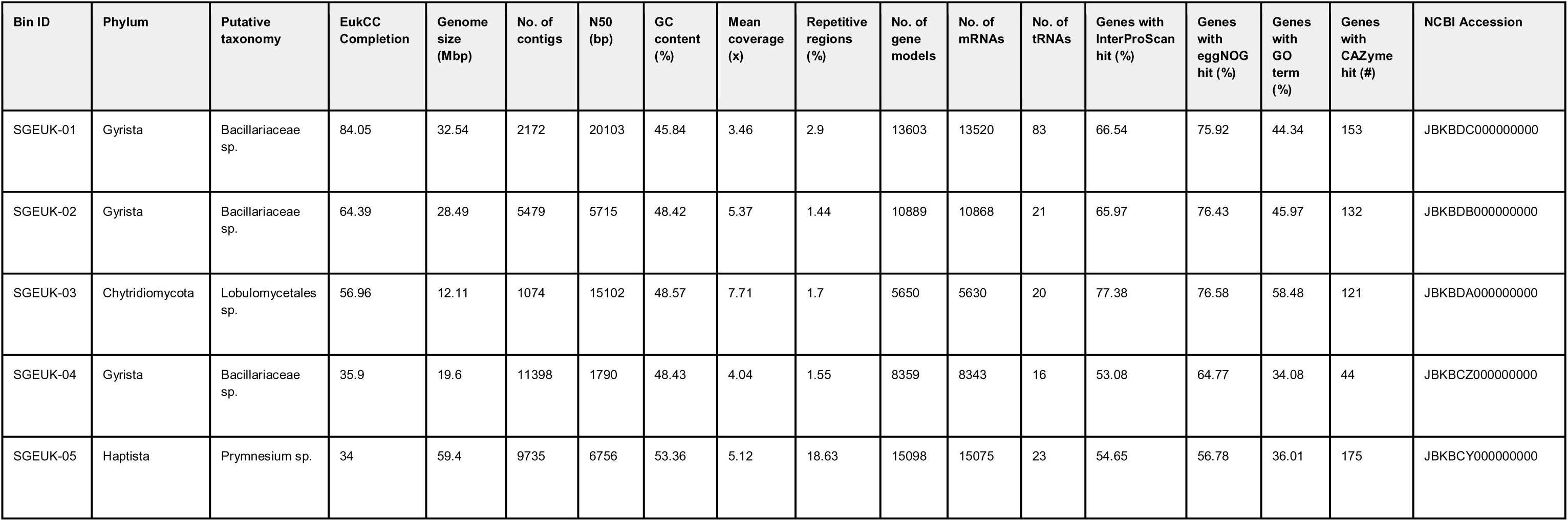
Eukaryotic metagenome-assembled genome assembly and annotation statistics. We identified five high-quality (> 30% complete) eukaryotic MAGs associated with *Z. marina* leaves. Here, we list their proposed taxonomy, report on various genome and annotation quality statistics, and provide the NCBI accession information for each MAG.

Overall, despite fragmentation, these five eukaryotic MAGs exhibited varying levels of genome and annotation completeness. The Bacillariophyta MAGs, particularly SG-EUK01 and SG-EUK02, displayed higher genome completeness, reflecting robust reconstructions of their genomic landscapes. SG-EUK03 showed a more moderate genome completeness, but a more robust annotation. SG-EUK04 and SG-EUK05 are more fragmented, though still represent valuable insight into the genomes and functions of these microbial eukaryotes.

### Phylogenetic placement of eukaryotic MAGs puts them in the diatom, fungal, and haptophyte trees of life

Following MAG recovery and functional annotation, we constructed phylogenetic trees to refine the taxonomic placement of the five draft MAGs. Using BUSCO protein sets and guided by preliminary taxonomic assignments from EUKulele, we built three trees: one for Bacillariophyta, one for fungi, and one for Haptista.

SGEUK-01, SGEUK-02, and SGEUK-04 were placed into the Bacillariophyta tree, and clustered among diatom lineages in the Bacillariaceae family, with SG-EUK02 and SG-EUK04 forming a clade that is sister to *Nitzschia inconspicua*, while SG-EUK01 formed a novel branch serving as outgroup to the clade representing all members of the Bacillariaceae family (Figure 1). While SGEUK-01 had very similar BUSCO completion estimates compared to annotated close relatives, both SGEUK-02 and SGEUK-04 likely paint a more incomplete picture of the functional landscape of these organisms. All three MAGs have genomes and annotations on the smaller side with publicly available genomes for members of the Bacillariaceae family ranging from 27.23 Mbp to 99.71 Mbp in size [123,214], and with available annotations ranging in gene count from 11,972 to 39,085 genes [123,139].

**Figure 1.**
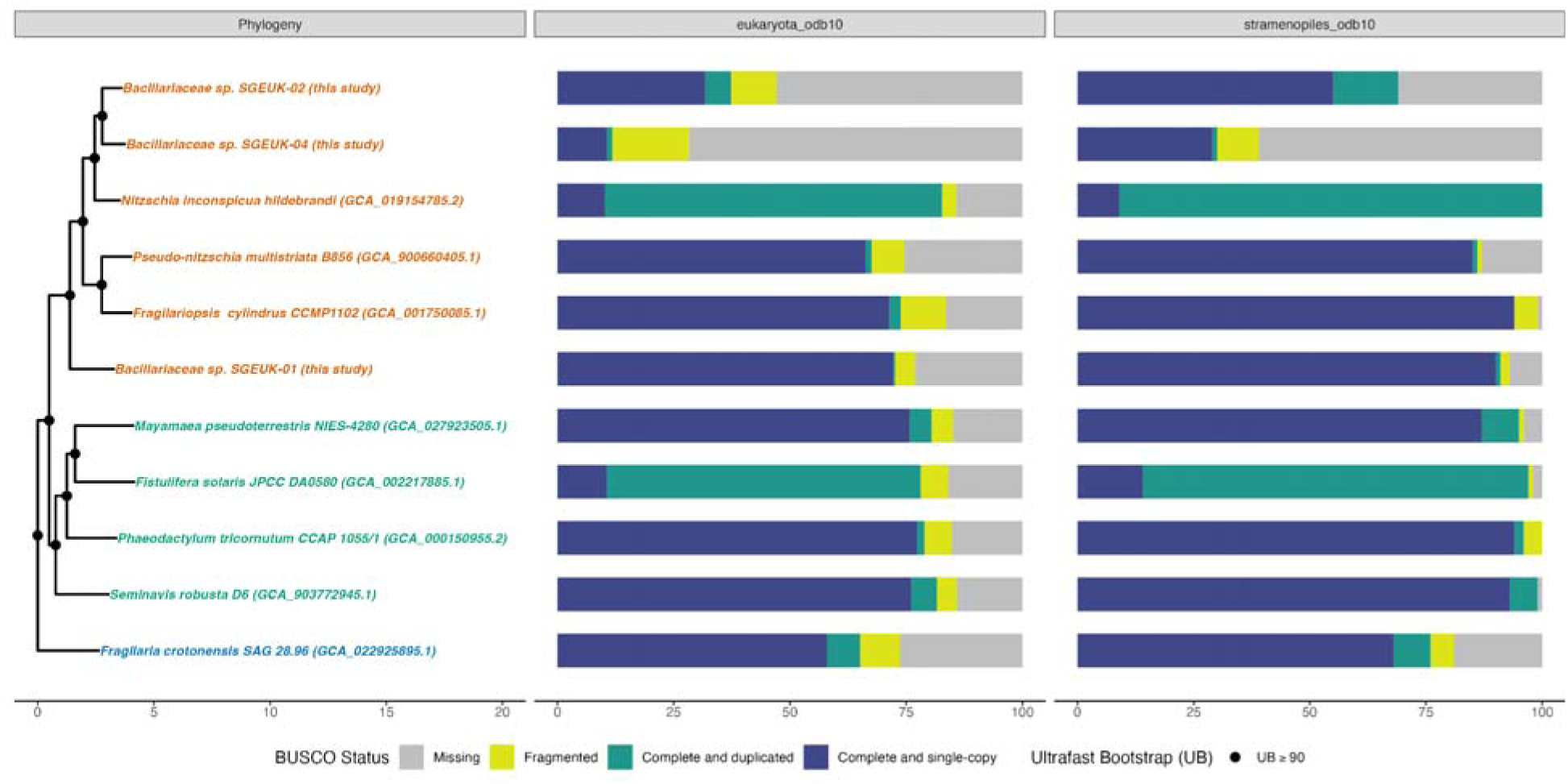
SGEUK-01, SGEUK-02 and SG-EUK04 are phylogenetically placed in the family Bacillariaceae. From left to right, first, a maximum likelihood phylogeny that shows the placement of SGEUK-01, SGEUK-02 and SG-EUK04 in the gyrista tree of life among diatom lineages. This tree was generated using IQ-TREE2 on an alignment of BUSCO stramenopiles_odb10 HMMs constructed using the PHYling_unified pipeline. Taxon labels in the phylogeny are shown colored by their assigned taxonomic family. Next, in association with this phylogeny, bar charts of the BUSCO “protein” completion status for the eukaryota_odb10 and stramenopiles_odb10 sets is shown. Bars show the percentage of genes found in each genome annotation as a percentage of the total gene set and are colored by BUSCO status (missing = gray, fragmented = yellow, complete and duplicated = green, complete and single copy = blue).

SGEUK-03 was placed into the fungal tree, falling amongst Chytridiomycota lineages (Figure 2). The MAG was placed sister to a clade containing *L. angularis* and *C. vesicula*, which both belong to the order Lobulomycetales, an undersequenced group of chytrids. *Quaeritorhiza haematococci*, a parasitic chytrid of algae [215], serves as an outgroup to the Lobulomycetales clade containing these three taxa. It falls in the same taxonomic class as Lobulomycetales, the Chytridiomycetes, but its phylogenetic placement in an order is uncertain. Regardless, it has been shown to be sister to the Lobulomycetales in other phylogenetic studies [216]. The two available Lobulomycetales genomes are similarly fragmented to SGEUK-03, with *L. angularis* represented by 1253 scaffolds with an N50 of 26,617 bp and *C. vesicula* spanning 2291 scaffolds with an N50 of 14,665 bp [172]. However, both their genomes (24.65-24.94 Mbp) and annotations (8737-9208 genes) are larger than SGEUK-03 which is in line with its lower BUSCO completion scores (Figure 2).

**Figure 2.**
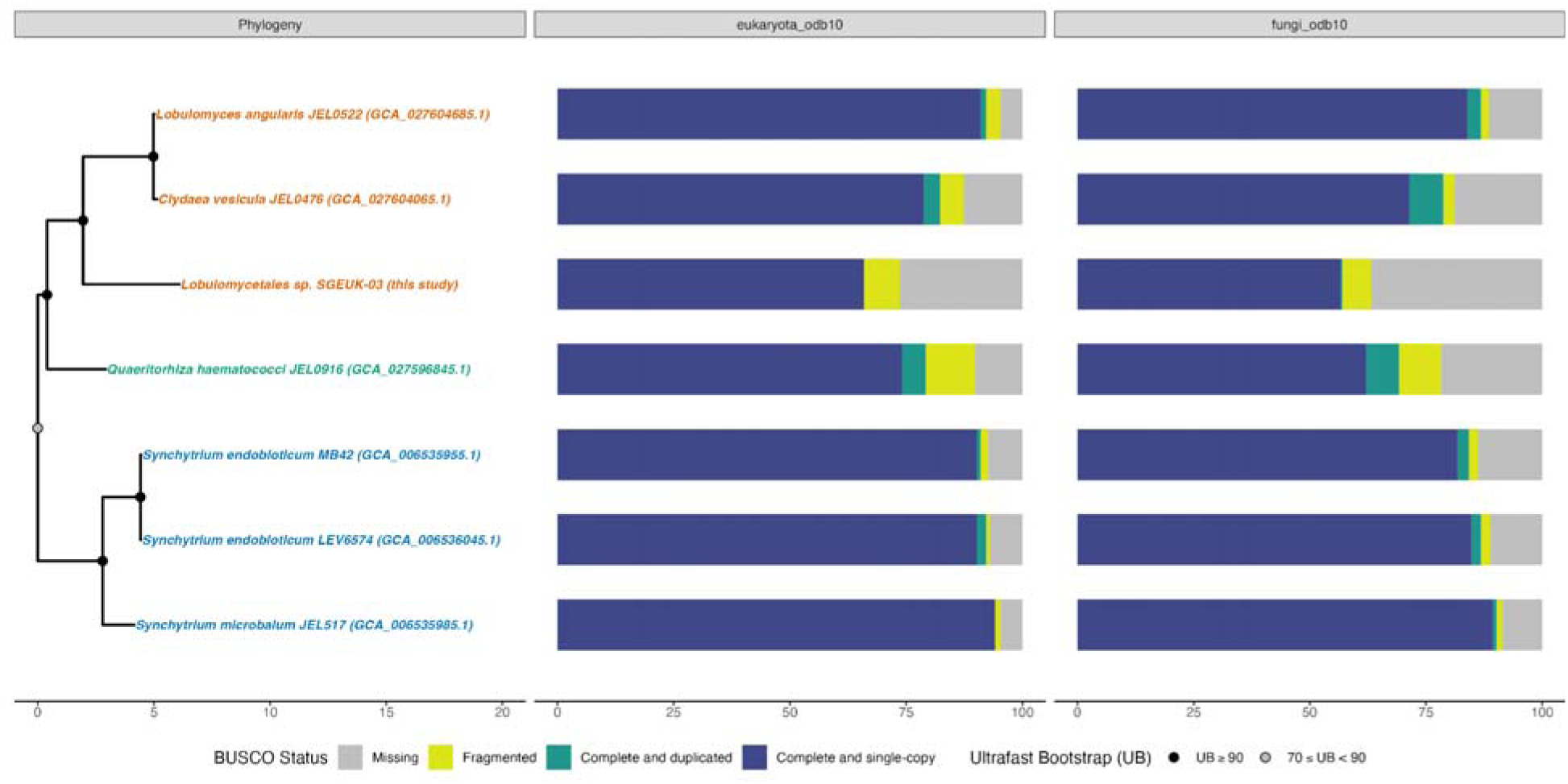
SGEUK-03 is phylogenetically placed in the order Lobulomycetales. From left to right, first, a maximum likelihood phylogeny that shows the placement of SGEUK-03 in the fungal tree of life. This tree was generated using IQ-TREE2 on an alignment of BUSCO fungi_odb10 HMMs constructed using the PHYling_unified pipeline. Taxon labels in the phylogeny are shown colored by their assigned taxonomic order. Next, in association with this phylogeny, bar charts of the BUSCO “protein” completion status for the eukaryota_odb10 and fungi_odb10 sets is shown. Bars show the percentage of genes found in each genome annotation as a percentage of the total gene set and are colored by BUSCO status (missing = gray, fragmented = yellow, complete and duplicated = green, complete and single copy = blue).

To further confirm and improve placement of SGEUK-03 among the Lobulumycetales, we used phyloFlash to recover an 18S rRNA gene from the metagenomic dataset likely belonging to this MAG and then placed in in the context of available 18S rRNA genes representing this chytrid lineage. This 18S rRNA gene phylogeny confirmed placement of SGEUK-03 among the Lobulumycetales, and further placed it within the novel SW-I clade, an aquatic clade of Lobuluymycetales largely described from DNA survey datasets (Figure 3). SGEUK-03 is sister to a sequence from deep sea marine sediment [199] and is in a clade with other sequences from marine surveys [198]. This taxonomic placement matches that previously obtained for the dominant amplicon sequence variant (SV8) from ITS2 amplicon profiling of the same samples, supporting that SGEUK-03 represents the chytrid fungus previously identified to be abundant on *Z. marina* leaves [36].

**Figure 3.**
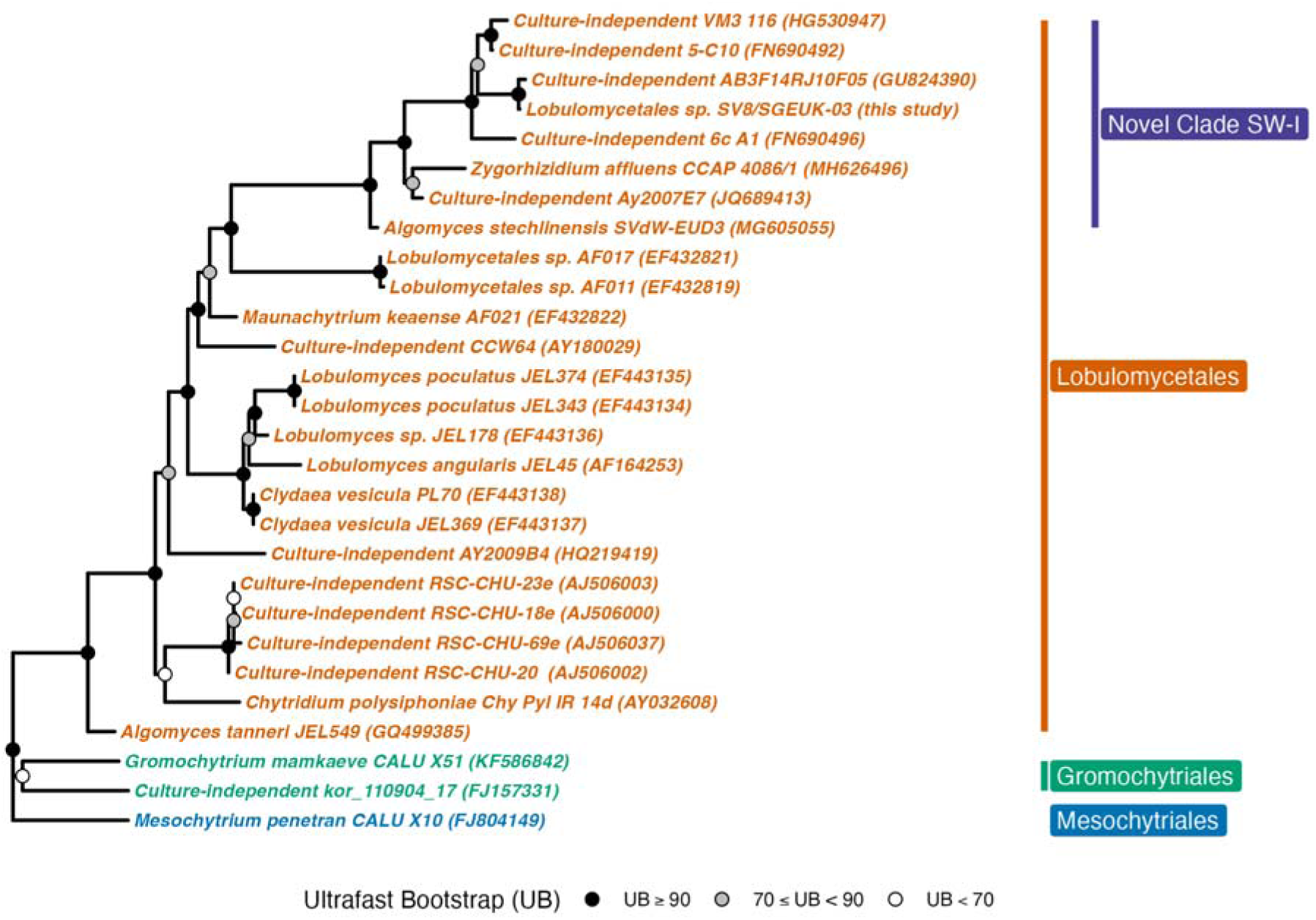
SGEUK-03 18S ribosomal RNA gene tree confirms placement in the Lobulomycetales in Novel Clade SW-I. Given the lack of closely related genomes to SGEUK-03, we used phyloFlash to assemble the 18S rRNA gene associated with SGEUK-03 and then aligned it to close relatives using SSU-align. The alignment was trimmed with trimAl, and IQ-TREE2 was run with ModelFinderPlus, to identify and use the optimal evolutionary model, and 1000 ultrafast bootstraps. Taxon labels in the phylogeny are colored and indicated by bars based on their assigned taxonomic order. Additionally, Novel Clade SW-I is also indicated by a bar. Displayed at each node as a circle in the tree are the ultrafast bootstrap values (e.g. a black circle represents bootstrap values greater or equal to 90%, a grey circle represents bootstrap values greater or equal to 70%, a white circle represents bootstrap values less than 70%).

SGEUK-05 was placed into the haptophyta tree, among the *Prymnesiales* order and sister to the transcriptome of the phytoplankton, *Prymnesium parvum* (Figure 4). SGEUK-05 and *P. parvum* were sister to a clade of transcriptomes from *Chrysochromulina polylepis* which are also members of the *Prymnesiaceae* family. *Chrysochromulina* genomes are 59.07-65.75 Mbp [189], though other members of the *Prymnesiales* have genomes up to ∼167.68 Mbp [187]. Given the relatively low completion of SGEUK-05, despite its larger size of 59.4 Mbp, it is likely we are only capturing less than half of its genomic and functional space.

**Figure 4.**
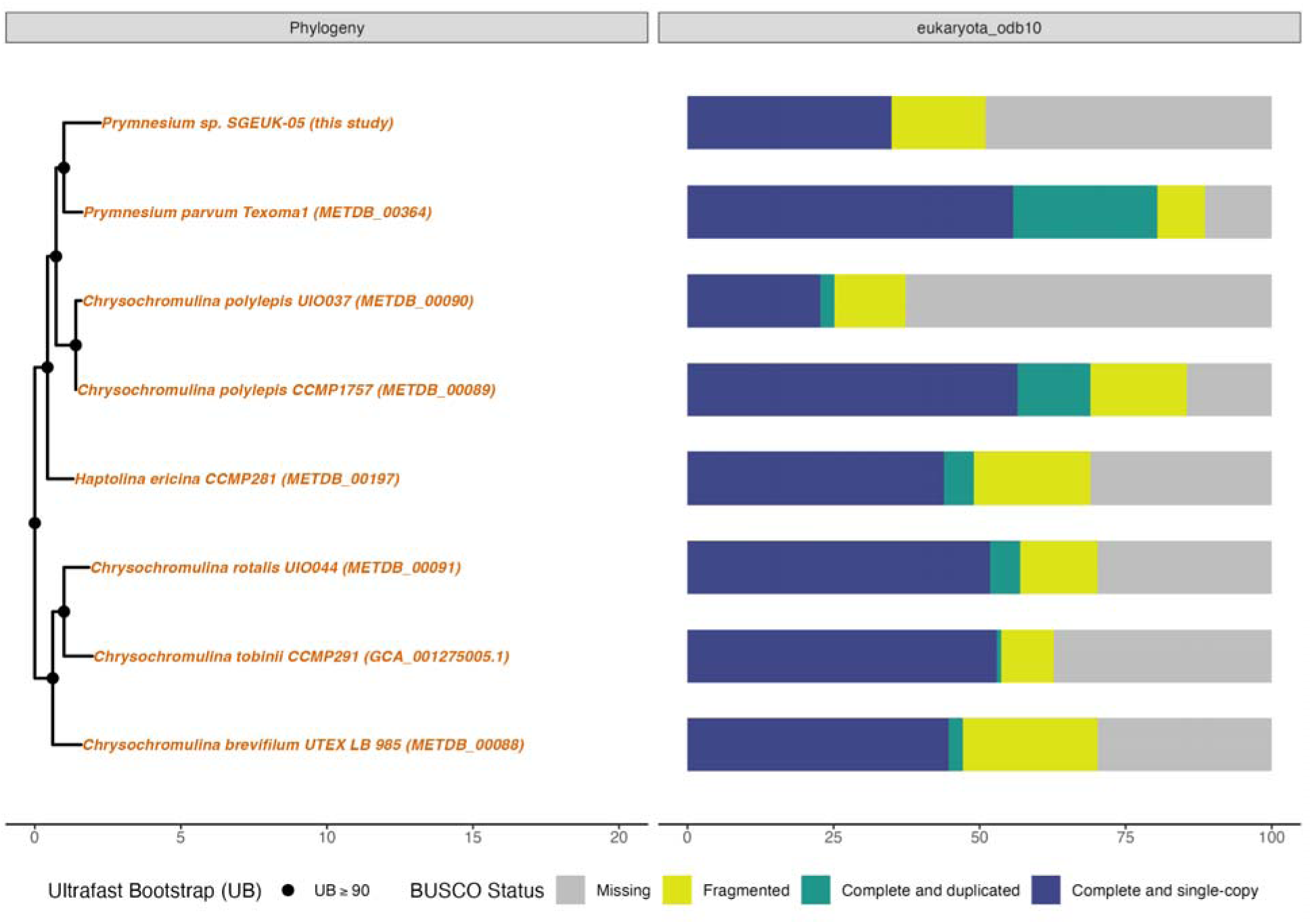
SGEUK-05 is phylogenetically placed in the genus Prymnesium. From left to right, first, a maximum likelihood phylogeny that shows the placement of SGEUK-05 in the haptista tree of life among haptophyte lineages. This tree was generated using IQ-TREE2 on an alignment of BUSCO stramenopiles_odb10 HMMs constructed using the PHYling_unified pipeline. Taxon labels in the phylogeny are shown colored by their assigned taxonomic family. Next, in association with this phylogeny, a bar chart of the BUSCO “protein” completion status for the eukaryota_odb10 set is shown. Bars show the percentage of genes found in each genome annotation as a percentage of the total gene set and are colored by BUSCO status (missing = gray, fragmented = yellow, complete and duplicated = green, complete and single copy = blue).

### Comparative genomics of SGEUK-03 reveals genes involved in host interaction

We performed OrthoFinder analysis, comparing SGEUK-03 with other fungi with a focus on comparisons to other members of the Lobulomycetales and Chytridiomycota. In total, we identified 75,458 phylogenetically hierarchical orthogroups (HOGs) among all fungi, of which 44,619 were identified among all Chytridiomycota. Of those, 8192 were identified among the Lobulomycetales, with a total of 4238 HOGs identified in SGEUK-03. Of the 5630 genes in SGEUK-03, 4886 genes (86.8%) were assigned to HOGs. Given that SGEUK-03 represents an incomplete MAG, we focused our analyses on high-copy number orthogroups (≥5 members) in SGEUK-03 and compared these to the average copy numbers in other fungal groups (Figure 5).

**Figure 5.**
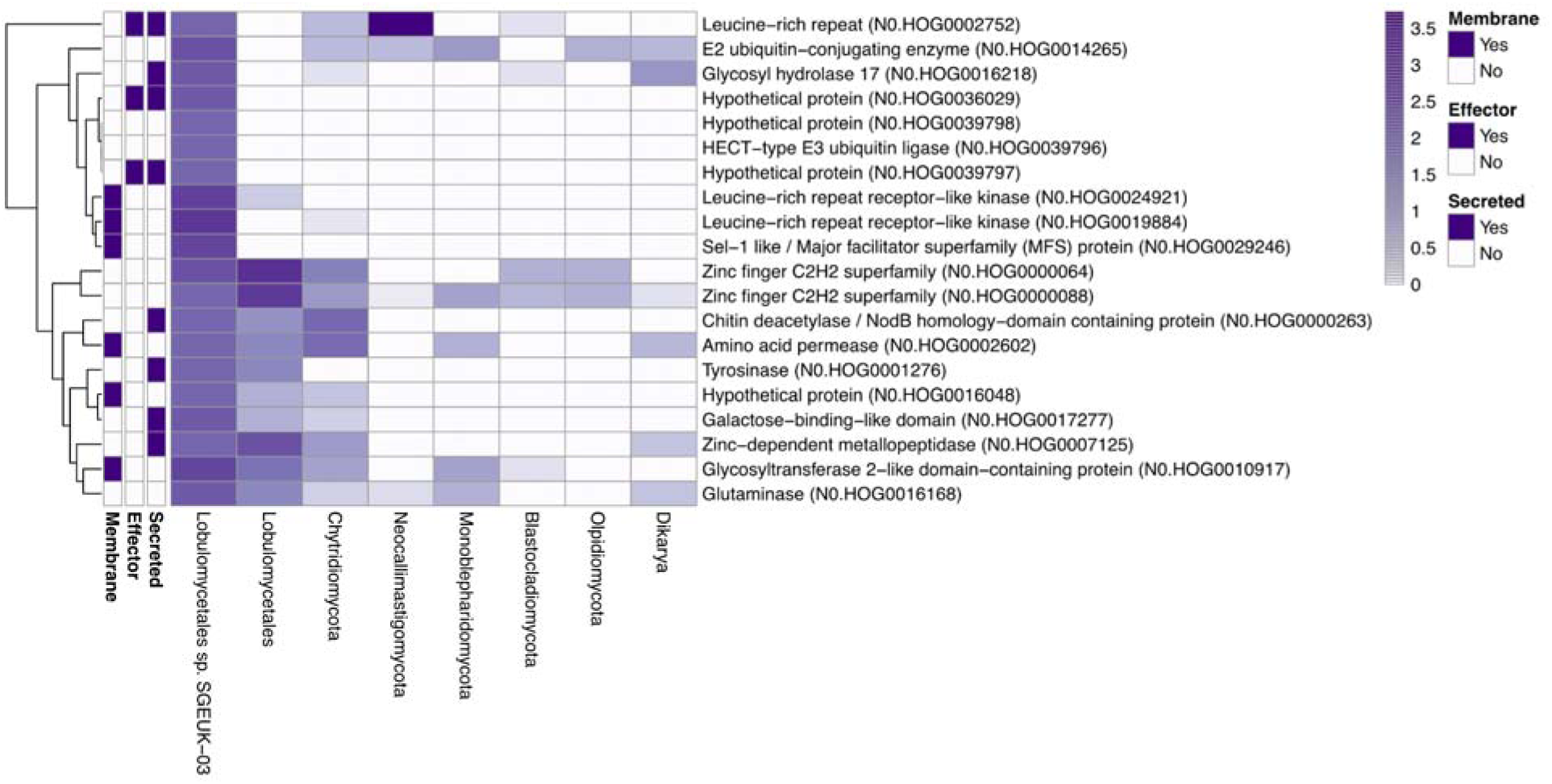
Only some high-copy orthogroups in SGEUK-3 are shared with close relatives. A heatmap is depicted here visualizing the log-scaled copy numbers of phylogenetically hierarchical orthogroups (HOGs) with 5 or more copies in SGEUK-03 in comparison to the average copy number of the same HOGs in other fungal taxonomic groups (i.e., other members of the order Lobulomycetales, other Chytridiomycota lineages, or other fungal phyla). Additionally, whether these genes are predicted in the annotation of SGEUK-03 to be membrane-bound, secreted or as plant-effector is indicated on the left of the heatmap. Further, the dendrogram to the left of the heatmap clusters the HOGs by similarity in log-scaled counts between the different HOGs.

In total there were 20 high-copy HOGs, of which five were unique to SGEUK-03. These included three hypothetical proteins (of which two were predicted cytoplasmic effectors), a Hect-type E3 ubiquitin ligase and a Sel-1 like / Major Facilitator Superfamily (MFS) protein. Six HOGs were shared exclusively with other Chytridiomycota or Lobulomycetales, including genes potentially involved in host interactions, such as leucine rich repeat receptor like kinases (LRR-RLKs), a galactose-binding like domain encoding gene, a tyrosinase, and a secreted chitin deacetylase (NodB homology-domain containing protein). Additionally, three HOGs were shared with other early-diverging fungal lineages, including a transcription factor, a GT2, and an LRR effector. Finally, six HOGs were shared across distant fungi, including Dikarya, such as GH17, transcription factors, and a glutaminase.

## Discussion

### Draft MAGs represent key ‘omics resources of under sequenced eukaryotic lineages

We report the successful recovery of five eukaryotic MAGs associated with the leaves of the seagrass *Z. marina*. The five draft MAGs represent under sequenced eukaryotic lineages including three MAGs representing diatoms in the family Bacillariaceae (93%, 70% and 31% BUSCO completeness), one MAG representing a haptophyte algae in the genus *Prymnesium* (40% BUSCO completeness), and a single fungal MAG in the order Loubulomycetales (65% BUSCO completeness). We annotated these five draft eukaryotic MAGs, providing an invaluable functional resource to enable foundational insight into their roles in the marine ecosystem. While all five MAGs represent relatively novel ‘omics resources, we highlight SGEUK-03 which we could only taxonomically place at the order level as part of the Loubulomycetales, a group with only two sequenced genomes, of which both are distant relatives of this MAG. Overall all five eukaryotic MAGs represent significant contributions of genomic novelty and we expect comparative genomic and functional approaches by the research community to further elucidate their evolution and ecology.

### Diatoms and Phytoplankton are critical, yet understudied, primary producers in seagrass ecosystems

Seagrass leaves serve as an ideal habitat and stable refuge for marine diatoms and algae. In fact, diatoms, including members in the Bacillariaceae, make up the biggest component of the microscopic epiphytic biomass associated with seagrass leaves [19,217–219], have the largest relative abundance within the microbial eukaryotic community associated with *Z. marina* globally [25] and are considered part of *Z. marina*’s core epiphytic leaf community [26,27]. Similarly, phytoplankton and other microalgae are another important part of the epiphytic community of seagrasses [18,24,220]. Both diatoms and microalgae make significant contributions to primary productivity, are seasonally variable and serve as important food sources in seagrass bed ecosystems [19,20,218,221–224]. In this work, seagrass leaves were free of a visible epiphytic biomass layer, suggesting that the recovery of these MAGs might indicate a rather tight association of these taxa with *Z. marina* leaf tissues. We hope these draft MAGs and their annotations provide useful resources for the community to begin to interrogate their genomics, possible functional roles, community dynamics in seagrass beds and the resilience of these microbial eukaryotes in the face of global climate change.

### Loubulomycetales may have a biotrophic or symbiotic role in seagrass ecosystems

SGEUK-03 clustered within Novel Clade SW-I in the Lobulumycetales (Figure 3). Known isolated members of Novel Clade SW-I are chytrids that are parasites of freshwater algae and diatoms [203,204,225]. While additional unclassified members of Novel Clade SW-I are only known from amplicon-based marine surveys of deep sea sediments, open ocean and seagrass ecosystems [36,198,199]. Overall, the Loubulomycetales are a relatively understudied group of chytrid fungi that have been found to span terrestrial, freshwater and marine ecosystems as both parasites and saprobes [43]. Given this diversity, and the absence of isolated representatives from seagrass environments, the ecological role of SGEUK-03 remains unclear. However, insights from its genome, specifically exploring effector and CAZyme profiles (Figure S1, Table S5), provide clues to its possible functional role.

Effectors are critical for all plant-fungal interactions [226–229] including for symbiotic and biotrophic fungi which still need to evade host plant recognition and suppress host defenses to persist [230,231]. Additionally, parasitic chytrids of animals and plants have been observed to have a repertoire of effectors proteins, as well as pathogenicity factors [166,170,232–234].

SGEUK-03 has a larger number of cytoplasmic effectors than apoplastic effectors, a pattern that might be consistent with an intracellular role. The majority of predicted effectors were hypothetical proteins with limited functional annotation. While the precise function of these effectors remains unclear, their abundance in SGEUK-03 points to a host-associated role.

We can explore the CAZyme diversity of SGEUK-03 to further hone in on its functional role. CAZyme-encoding genes vary among fungal species and likely reflect their adaptation to different nutritional niches [235]. For example, plant pathogens have been found to have higher numbers of CAZymes than saprophytic and animal pathogens [236]. Plant cell wall degrading enzymes, such as GHs, CEs and PLs, in particular, are often employed by fungal plant pathogens [237]. Necrotrophic pathogens deploy these cell wall-degrading enzymes to promote host damage and colonization [238]. Whereas endophytic fungi and biotrophs tend to have fewer of these cell wall degrading enzymes [238–241]. While CAZyme diversity has been widely explored on land, particularly in the Dikaryan fungi, little characterization of these enzymes has been done in chytrids, though initial reports suggest these fungi can have a wide range of CAZymes [242].

SGEUK-03 had relatively few CAZymes overall, with a notable absence of enzymes reported to be involved in the degradation of plant cell wall components such as cellulose (GH6, GH7, GH12, GH45, GH3, GH61, CBM1), hemicellulose (GH10, GH11, GH74, GH27, GH36, GH26, GH43, GH51, GH54, GH62, GH35, GH67, GH115), and pectin (GH28, GH78, PL1, PL3, PL4, PL9) [236,239,243,244]. Instead, the glycosyl transferases (GTs) were the most represented group (53.7% of CAZymes), with GT2 being the most frequent, a pattern perhaps consistent with fungi that modify their own cell wall to evade detection, rather than degrade host cell walls.

Interestingly, the CAZyme profile of SGEUK-03 mirrors that of the obligate mutualistic symbiont *Rhizophagus irregularis*, an arbuscular mycorrhizal fungus that engages in a mutualistic relationship with land plants [245]. The relatively low CAZyme diversity of SGEUK-03, combined with its effector content, suggest that it may engage in a similar strategy of minimal host damage while exploiting host resources. This idea is further supported by the prediction from CATAStrophy [98], which classified SGEUK-03 as a symbiont based on its CAZyme content. However, it is important to note that CATAStrophy was trained on filamentous dikaryon fungi, and its application to chytrids may be limited. Additionally, as the genome of SGEUK-03 is incomplete, we cannot rule out the presence of additional CAZymes that could alter this classification. Regardless, on the whole, the effector and CAZyme profiles of SGEUK-03 may suggest that it has a symbiotic or biotrophic role in seagrass ecosystems.

### Are Lobulomycetales chytrids biotrophic parasites or symbionts of seagrasses themselves?

The frequent identification of *Lobulomycetales* species in *Z. marina* leaf epiphyte and endophyte samples from previous amplicon surveys [25,36] supports the hypothesis that SGEUK-03 may be closely associated with *Z. marina*. Microscopic observations of chytrids directly interacting with seagrass tissues further support a possible symbiotic or weakly parasitic role [246]. However, given the diversity of hosts that chytrids can associate with, we cannot rule out the possibility that SGEUK-03 may also be parasitizing a diatom or algal host that is itself associated with *Z. marina*. While we saw no clear coverage pattern (e.g., 1-to-1) which might suggest one of the other MAGs might be a host for SGEUK-03, another possibility could be that SGEUK-03 is a generalist chytrid, capable of associating with multiple hosts including *Z. marina* and members of its epiphytic community.

Chytrids are well known parasites of a broad range of phytoplankton, including diatoms, green algae, dinoflagellates, and cyanobacteria and can display both high host specificity or broad generalist strategies, depending on environmental conditions and host availability [203,204,247–255]. A generalist strategy allows for infection of multiple species enabling chytrids to persist despite fluctuations in available host communities, with trade-offs including lower growth rates as a result of needing to maintain the genomic potential to evade diverse host immune responses [251]. Members of the Lobulomycetales and marine fungi more generally have been observed to exhibit some seasonality in their emergence which has been hypothesized to relate to host-dynamics and environmental conditions [44,256–258]. Blooms of parasitic chytrids, for example, frequently accompany an increased abundance of diatoms that the chytrids infect and ultimately kill. Previous work has suggested that Lobulomycetales sp. associated with seagrasses experience bloom-like dynamics, which could relate to seasonality, environmental conditions or host availability [25,36]. It is possible that the presence of SGEUK-03 in seagrass ecosystems is similarly influenced by seasonal patterns in host availability, whether that host is seagrass itself or associated epiphytic algae or diatoms. Further work is needed to confirm and explore the host-specificity of SGEUK-03.

### SGEUK-03’s high copy orthologs may represent a functional repertoire focused on host association

To better understand its functional potential, particularly its interactions with its host, we used comparative genomics approaches to examine SGEUK-03’s highest copy orthologs. We found twenty high-copy gene families with functions that indicate they may be involved in host or environmental recognition, evasion of host immune responses, and nutrient acquisition.

Among the high-copy orthogroups, we identified two LRR-LRKs, which are receptor kinases that detect external signals and are often implicated in non-self recognition. For example, LRR-LRKs are best known for the role they play in plant and animal innate immunity [259,260]. Dramatic gene family expansions of LRR-LRKs are observed in plants and animals, but these genes are largely missing in dikaryon fungi, whose innate immune system is still not fully understood, though a small number of LRR-LRKs have been reported in chytrids and zygomycetes [261–263]. The HOGs that included LRR-LRKs were shared with other chytrids and were the most abundant HOGs identified for SGEUK-03. Aquatic chytrids have motile zoospores which enable the targeting of specific hosts and substrates, a capability that is absent in dikaryan spores which lack a flagella [45]. For instance, chytrid zoospores can be attracted to dissolved molecules in the environment which may facilitate host detection [264–266]. We speculate that these LRR-LRKs may be involved in environmental sensing and host detection in SGEUK-03, although a role in innate immunity, similar to that of plants and animals, could also be possible.

A galactose-binding domain protein, another high-copy gene family shared with chytrids, may support early host recognition and adhesion. Galactose is an important sugar in seagrass cell walls [267]. Additionally, a galactose-binding CBM plays an important role in host-fungal interactions through mediating chemotaxis towards the host, attachment, and virulence in a chytrid pathogen [268]. SGEUK-03 may be similarly using this gene family for sensing and adhering to host tissues during colonization, establishing an interaction with the host early in the infection process.

In addition to host recognition, SGEUK-03 may possess mechanisms to evade host immune responses during and after invasion. Fungi have evolved several strategies to evade host immune responses including modifying their cell wall composition and secreting effector proteins to protect cell walls or help evade host detection [269,270]. SGEUK-03 possesses a suite of effectors, including two unique secreted cytoplasmic effector families. In addition, SGEUK-03 encodes a unique family of Hect-type E3 ubiquitin ligases. These ligases play critical roles in regulating protein degradation through modification, which impacts cellular processes such as stress responses, endocytosis, cell cycling, and pathogenicity [271–275] Such effectors and ubiquitin ligases may provide SGEUK-03 with the ability to manipulate host cell machinery and suppress immune responses.

Additionally, SGEUK-03 shares a gene family annotated as a chitin deacetylase (NodB homology-domain containing protein) with other chytrids. Chitin deacetylases can modify the fungal cell wall during plant invasion making it less recognizable to plant defense systems [276,277]. Chitin deacetylases with NodB homology are also involved in invasion and signaling by symbiotic fungi [278]. SGEUK-03 may similarly modify its cell wall to evade host immunity or maintain host interactions.

SGEUK-03 has a glycoside hydrolase family 17 (GH17) enzyme gene family, which it shared with distant fungal lineages, including members of Dikarya. GH17 enzymes are involved in the modification of cell wall glucan in both fungal and plant cells, which is essential for the growth, development and host-pathogen interactions [279–281]. On land, *Cladosporium fulvum*, a hemibiotrophic fungal pathogen of tomatoes, secretes a GH17 which triggers release of a damage-associated molecular pattern from the host cell wall triggers host innate immunity and ultimately cell death [282]. Further, GH17 enzymes in fungi are abundant and thought to have important roles in global carbon cycling in the open ocean [283]. For example, a marine *Cladosporium* species secretes GH17 when utilizing laminarin [284], which is an algal and diatom-derived β-glucan that’s abundant in the ocean [285]. It is possible that GH17 enzymes in SGEUK-03 could play a similar role in interacting with host cell walls, either during invasion or as part of a defensive strategy.

Tyrosinases, a high-copy gene family of SGEUK-03 shared with other chytrid lineages, have been previously reported as expanded and implicated in host-pathogen interactions [286,287]. These enzymes are involved in melanin biosynthesis, which contribute to fungal virulence by protecting cells from UV radiation, reactive oxygen species (ROS), and hydrolytic enzymes that degrade fungal cell walls [288–290]. Melanin production could provide SGEUK-03 with protection from host defenses, as ROS accumulation is a common plant response to pathogen recognition [238,291]. Therefore, the putative tyrosinases in SGEUK-03 might provide protection from ROS-mediated host defenses, ensuring its survival within host tissues.

We also found that gene families with functions likely related to nutrient acquisition, detoxification and stress responses including several transcription factors and transporter genes. This included a unique Sel-1 like / MFS transporter family, which are a group of transporters vital in cross-membrane transport of organic solutes. They are key in fungal nutrient acquisition and may play roles in detoxification or efflux of harmful plant compounds [292]. Overall, the observed suite of high-copy orthologs in SGEUK-03 supports a functional landscape of gene families with putative annotations related to host or environmental recognition, evasion, and interaction.

## Conclusion

We successfully recovered five eukaryotic MAGs from DNA isolated from the leaves of the seagrass *Zostera marina*, representing previously under-sequenced eukaryotic lineages. These include three MAGs of diatoms from the family Bacillariaceae, one MAG from a haptophyte algae in the genus *Prymnesium*, and one fungal MAG from the order Loubulomycetales. The functional annotation of these draft MAGs offers a valuable resource for understanding their ecological roles in the marine environment. Notably, we placed SGEUK-03 within the novel clade SW-I of the Loubulomycetales and performed comparative genomic analysis, suggesting a symbiotic or biotrophic lifestyle based on its effector and CAZyme repertoire. Further exploration of high-copy orthogroups in SGEUK-03 revealed a suite of gene families potentially involved in host or environmental interactions, recognition, and immune evasion. While additional research is required to clarify host specificity and further characterize these MAGs, our findings contribute to the growing knowledge of marine microbial eukaryotic diversity and provide a genomic foundation for future studies on the ecological roles and evolutionary trajectories of these enigmatic microorganisms.

## DNA Deposition

The raw metagenomic sequencing data was deposited at GenBank under accession no. <u>PRJNA1140276</u>. MAGs and annotations under this BioProject at DDBJ/ENA/GenBank under accession no. JBKBCY000000000-JBKBDC000000000. The 18S rRNA gene obtained for SGEUK-03 was deposited at GenBank under accession no. PQ146498. All code used in this work has been deposited on Github (<u>casett/ZM_Euk_MAGs</u>) and archived in Zenodo (DOI: <u>10.5281/zenodo.14278507</u>).

## COI

JAE is on the Scientific Advisory Board of Zymo Research, Inc. JES is a scientific consultant for Michroma, Inc.

## Funding sources

The sequencing data in this work was generated by grants from the UC Davis H. A. Lewin Family Fellowship and the UC Davis Center for Population Biology to CLE. CLE was supported by the National Science Foundation (NSF) under a NSF Ocean Sciences Postdoctoral Fellowship (Award No. 2205744). JES is a CIFAR Fellow in the program Fungal Kingdom: Threats and Opportunities and was partially supported by NSF awards EF-2125066 and IOS-2134912. Computations were performed using the computer clusters and data storage resources of the UC Riverside HPCC, which were funded by grants from NSF (MRI-2215705, MRI-1429826) and NIH (1S10OD016290-01A1). The funders had no role in study design, data collection and analysis, decision to publish, or preparation of the manuscript.

## Author Contributions

CLE conceived and designed the experiments, performed sampling, analyzed the data, prepared figures and/or tables, wrote and reviewed drafts of the paper. JAE and JES reviewed drafts of the paper.

## Acknowledgements

We would like to thank Katherine Dynarski (<u>ORCID: 0000-0001-5101-9666</u>) and Sonia Ghose (<u>ORCID: 0000-0001-5667-6876</u>) for their help with initial sample collection. We would also like to thank John J. Stachowicz for use of his scientific sampling permit, California Department of Fish and Wildlife Scientific Collecting Permit # SC 4874.

## Notes

### Competing Interest Statement

Jonathan A. Eisen is on the Scientific Advisory Board of Zymo Research, Inc. Jason E. Stajich is a scientific consultant for Michroma, Inc.

### Summary of Updates

Corresponding author email updated; no changes to manuscript

