## Supplemental Figures and Table Legends for "Eukaryotic MAGs recovered from deep metagenomic sequencing of the seagrass, *Zostera marina,* include a novel chytrid in the order Lobulomycetales"

**Supplemental Table Legends and Figures:**

**Table S1.** Genomes used in phylogenetic analysis of gyrista. Here we provide details on each genome obtained from NCBI including assembly statistics, BioProject number, assembly accession number, and appropriate data citation.

**Table S2.** Genomes used in phylogenetic and orthogroup analyses of fungi. Here we provide details on each genome obtained from NCBI including taxonomic information, assembly statistics, BioProject number, and assembly accession number. We also report the predicted trophic mode of each organism and appropriate data citation.

**Table S3.** Genomes used in phylogenetic analysis of haptista. Here we provide details on each genome obtained from NCBI or METdb including BUSCO statistics, assembly accession number, and appropriate data citation.

**Table S4.** Sequences used in phylogenetic analysis of SGEUK-03 18S ribosomal RNA gene. Here we provide details on each sequence obtained from NCBI including taxonomic clade, species and strain ID, as well as NCBI accession number, and appropriate data citation.

**Table S5.** CAZyme distribution of SGEUK-03. Here we provide the counts of annotated genes with CAZyme domain hits, including the CAZyme category, CAZyme domain and gene count.

**Figure S1.** Effector and CAZyme profiles for SGEUK-03. (A) Bar plot depicting the number of secreted genes in the SGEUK-03 genome that are predicted to be plant apoplastic and cytoplasmic effectors. (B) Bar plot depicting the frequency distribution of predicted CAZymes in the SGEUK-03 genome across CAZyme families.

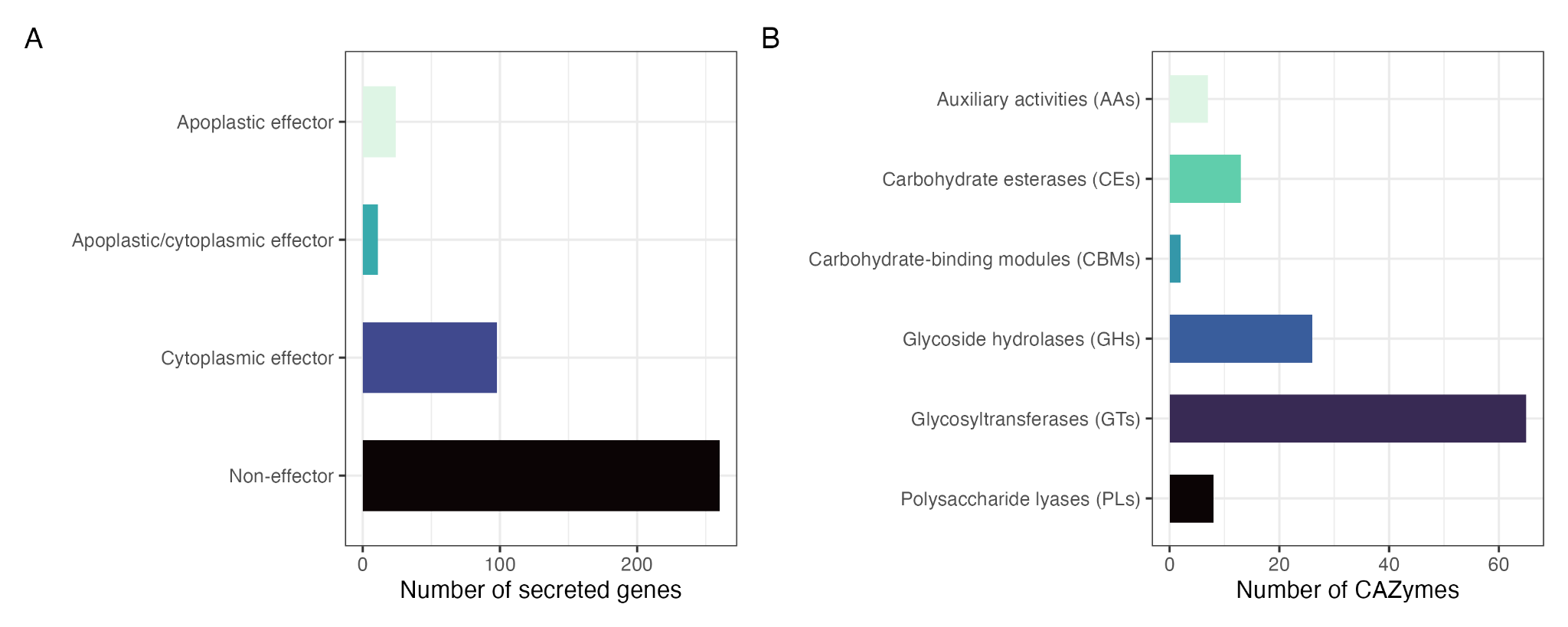
